# Endosomes facilitate mitochondrial clearance by enhancing Parkin recruitment to mitochondria

**DOI:** 10.1101/2020.02.19.955880

**Authors:** Rishith Ravindran, Anoop Kumar G. Velikkakath, Nikhil Dev Narendradev, Aneesh Chandrasekharan, T. R. Santhoshkumar, Srinivasa M. Srinivasula

**Affiliations:** School of Biology, Indian Institute of Science Education and Research Thiruvananthapuram, Thiruvananthapuram - 695551, Kerala, India; Cancer Research Program-1, Rajiv Gandhi Centre for Biotechnology, Thiruvananthapuram - 695014, Kerala, India

**Author notes:** These authors contributed equally to this work (Rishith Ravindran, Anoop Kumar G. Velikkakath). Correspondence to Srinivasa M. Srinivasula.

**Keywords:** CARP2, Endocytic vesicles, Mitophagy, PARK2, Ubiquitin ligase

## Abstract

Mutations in ubiquitin ligase Parkin are associated with Parkinson’s disease and defective mitophagy. Conceptually, Parkin-dependent mitophagy is classified into two phases; 1. Parkin recruits to and ubiquitinates mitochondrial proteins, 2. Formation of autophagosome membrane, sequestering mitochondria for degradation. Recently, endosomal machineries were reported to contribute to the later stage for membrane assembly. We report a role for endosomes in the events upstream of phase 1. We demonstrate that an endosomal ubiquitin ligase CARP2 associates with damaged mitochondria, and this association precedes that of Parkin. CARP2 interacts with Parkin, and stable recruitment of Parkin to damaged mitochondria was substantially reduced in CARP2 KO cells. Our study unravels a novel role of endosomes in modulating upstream pathways of Parkin-dependent mitophagy initiation.

## Introduction

Stringent quality control mechanisms ensure organelle and cellular homeostasis. Signaling pathways that regulate mitochondrial quality and function are believed to play critical roles in diverse physiological processes including cell survival and their dysregulation results in the development of numerous pathological conditions like neurological disorders, cancer and myopathies [1–3]. Autophagy is a conserved intracellular catabolic process in which cytoplasmic contents, including damaged mitochondria, are sequestered in double-membrane autophagosomes and delivered to the lysosomal compartment for degradation [4]. Selective elimination of dysfunctional and damaged mitochondria involving autophagic machinery and lysosomes is known as mitophagy [5].

Though cells can employ various other cellular tools like ESCRT machinery [6], MDVs (Mitochondrial Derived Vesicles) [7, 8] etc., to remove mitochondrial contents, mitophagy mediated by PINK1/Parkin is relatively well studied [9–11]. In depolarized mitochondria, PINK1 accumulated on the outer mitochondrial membrane phosphorylates Parkin, which in turn pleiotropically ubiquitinates several mitochondrial proteins result in the clearance of mitochondria by mitophagy [12]. However, details of the sequence of events involved in the Parkin recruitment, stability and identities of the proteins functioning in this pathway are not clear.

However, recent reports suggested a role for endosomes and endosomal proteins in mitochondrial clearance. Crosstalk of endosomal related proteins RABGEF1, Rab5, MON1/CCZ1 complex, and Rab7A result in Atg9A vesicle recruitment to autophagosome formation site [13]. Moreover, Rab5-positive endosomes have been implicated in the elimination of damaged mitochondria via the ESCRT complex machinery, and contribute to cytoprotective roles against oxidative stress [6, 14]. It has also been proposed that Rab11A endosomes enable the autophagy of damaged mitochondria [15]. How endosomes influence the mitochondrial quality and the crosstalk between the endosomal machinery and the above molecules that are known to be critical for mitophagy, remain to be elucidated.

CARP2 (Caspase-8- and -10-associated RING Protein 2) is one of the two RING and FYVE-like domain-containing proteins in the human genome. While the RING domain confers ubiquitin ligase activity, the FYVE-like motif along with its N-terminal sequence is reported to associate with phospholipids like phosphatidylinositol 3-phosphate [16]. Interestingly, CARP2-positive vesicles are also reported to be positive for Rab5, Rab7, Rab9, Rab11, and Lamp1 markers of diverse endocytic vesicles [17–19]. In this study, we report that CARP2-positive vesicles associate with damaged mitochondria and contribute to mitochondrial elimination by facilitating Parkin recruitment.

## Results

### Endosomal ubiquitin ligase CARP2 associates with damaged mitochondria

Because the endosomal machinery has been reported to contribute to mitophagy, we investigated the role of CARP2-positive endosomes in mitochondrial elimination. For these studies, we have used cells stably expressing CARP2 cloned under a weak promoter using a retroviral vector [20]. As a first step, we monitored the association between CARP2 endosomes and damaged mitochondria using confocal microscopy of A549 lung carcinoma cells stably expressing CARP2-EGFP. In these cells, mitochondria were stained with MitoTracker Red CMXRos. As expected, in cells treated with DMSO, CARP2-EGFP was primarily localized to intracellular vesicles, which were reported to be positive for various known endosomal markers (Figure 1A, [19]). In these cells, mitochondria appeared mostly as filamentous structures, with a minimum association with CARP2 vesicles (Figure 1C). However, mitochondria in cells treated with the mitochondrial uncoupler CCCP were found to be fragmented as expected, with substantial numbers being surrounded by CARP2 vesicles (Figure 1A). Line scan analysis demonstrated clear encircling of damaged mitochondria by CARP2 (Figure 1B), with nearly 60% of the live cells showing a positive association between these organelles within 25min of treatment (Figure 1C). Pearson coefficient analysis also showed a positive correlation in CCCP treated cells compared with untreated cells (Figure 1D). Another mitochondrial uncoupler commonly used is valinomycin. Results from cells treated with valinomycin also showed a similar association with what was observed in cells treated with CCCP (Figure S1A).

**Figure 1.**
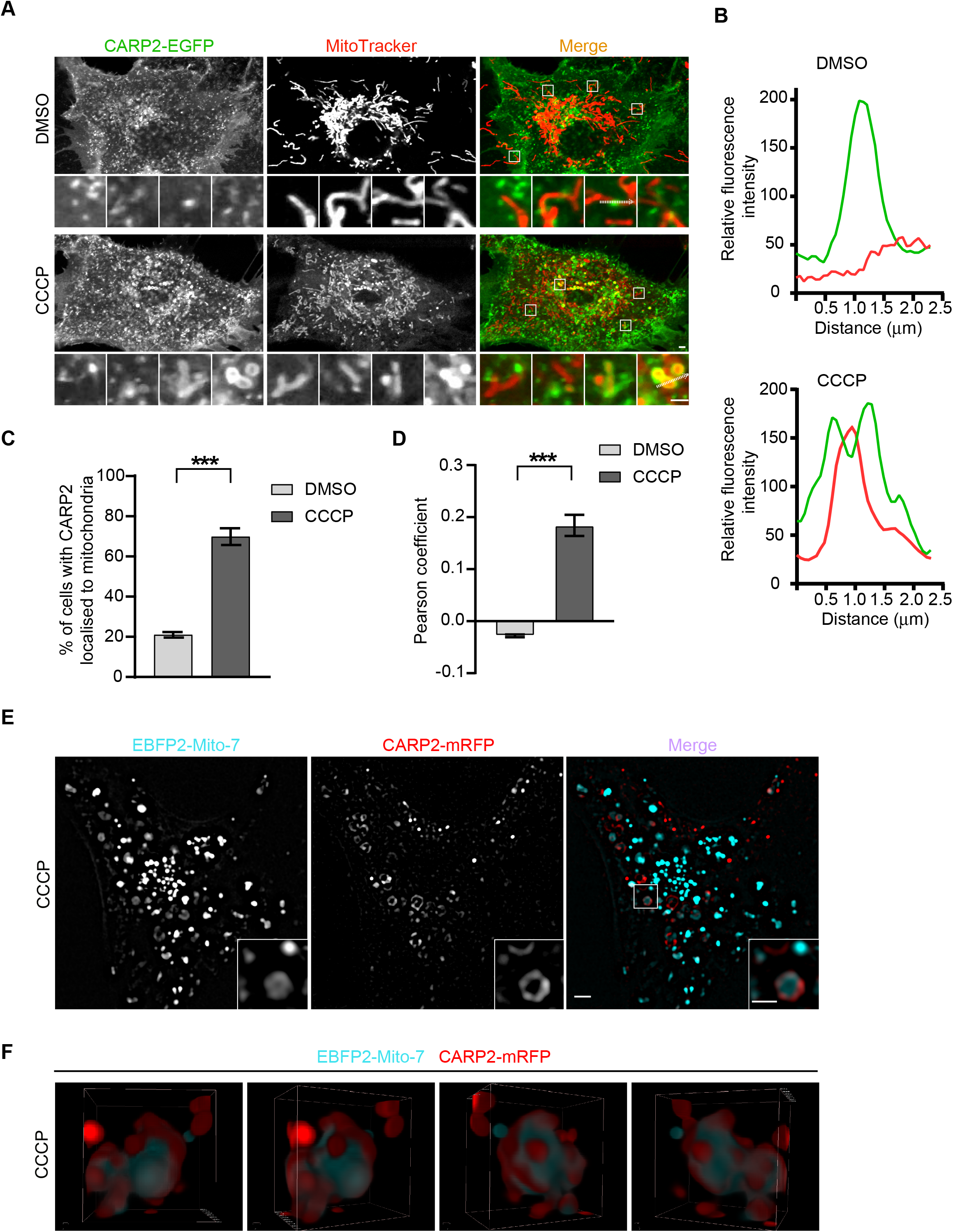
Endosomal ubiquitin ligase CARP2 associates with damaged mitochondria. (**A**) A549 cells stably expressing CARP2-EGFP stained with MitoTracker Red CMXRos for 30min, were washed with DMEM followed by treatment with DMSO or 20μM CCCP. Images of live cells were taken during 10-25min of CCCP treatment. Scale 2μm, inset scale 1μm. (**B**) Line scan analyses of the area under the arrow are shown. (**C**) Percentage of cell with CARP2-EGFP-positive mitochondria (MitoTracker Red CMXRos) was quantified using manual counting (cells having more than four CARP2 localized on MitoTracker were considered for counting). (**D**) The Association of CARP2 with mitochondria is calculated using Pearson correlation coefficient. Error bars represent mean ± SEM in cases of (**C**) and (**D)** from three independent experiments with a minimum of 50 cells per experiment. Statistical significance was calculated using a two-tailed unpaired t-test. P-values are <0.0001 in the case of **C** and **D**. (**E**) U2OS cell line stably expressing EBFP2-Mito-7 and CARP2-mRFP along with YFP-Parkin (not shown) were treated with 20μM CCCP and images of live cells were captured using super-resolution microscopy at 14 h post-treatment. Scale 2μm, inset scale 1μm. (**F**) Super-resolution 3D still images in four different angles of EBFP2-Mito-7-positive mitochondria and CARP2-mRFP-positive endosomes shown as a movie (**Movie 1**).

The association of CARP2 vesicles with damaged mitochondria was further analyzed by super-resolution imaging [Structured Illumination Microscopy (SIM)] in cells stably expressing EBFP2-Mito-7, CARP2-mRFP and YFP-Parkin (Figure 1E). The encirclement of damaged mitochondria by CARP2 positive vesicles was also noted in 3D super-resolution SIM images (Figure 1F and Movie 1).

To understand whether the association between CARP2 and mitochondria occurs cell-type independent manner, we have explored the association of CARP2 with mitochondria using neuronal cell line Neuro-2a (N2a), as mitochondrial quality maintenance is known to be critical for neuronal cell homeostasis [21, 22]. For this, we generated N2a cells stably expressing CARP2-mRFP. As it was noted in A549 cells, CARP2 appeared to associate with intracellular vesicles in N2a cells as well (Figure 2A, B, and S1B). In N2a cells also very little (basal level) association was noted between CARP2 and mitochondria (Mitotracker Deep Red FM) under DMSO treated conditions. However, as observed in CCCP treated A549 cells, a significant colocalization between CARP2-mRFP and damaged mitochondria was noted in N2a cells (Figure S1B, C). A similar association was also observed in other cell lines like human embryonic kidney cells, HEK 293T (data not shown) and osteosarcoma cell line, U2OS (Figure 3C), suggesting that CARP2 association with damaged mitochondria is cell-type independent.

**Figure 2.**
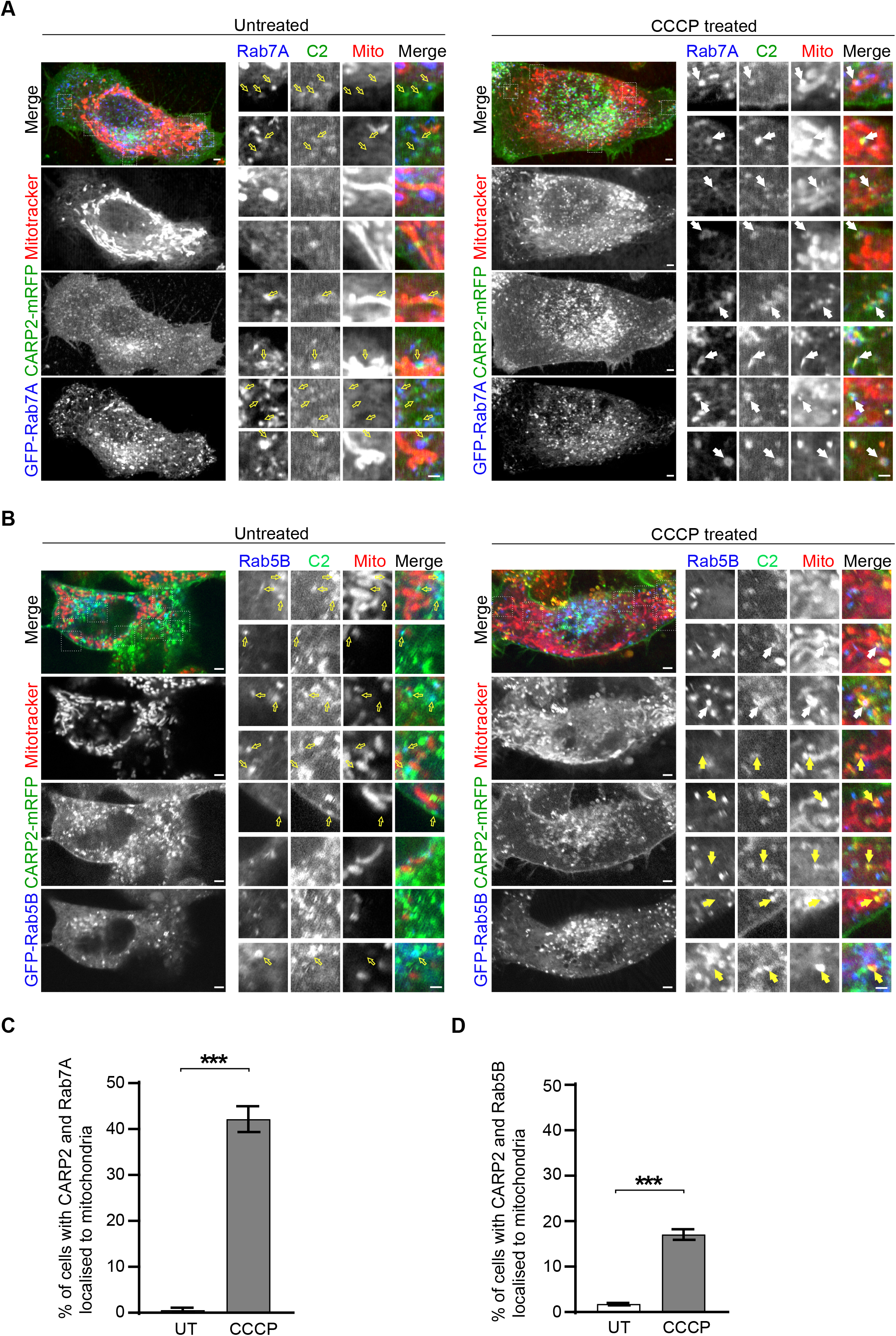
CARP2-associated mitochondria are positive for Rab7A and Rab5B. (**A** and **B**) Neuro-2a cells stably expressing CARP2-mRFP were transiently transfected with either GFP-Rab7A (**A**) or GFP-Rab5B (**B**). After 18 h of transfection, cells were stained with MitoTracker Deep Red FM for 30min and were subsequently treated with 10μM CCCP. Images of live cells from 10 to 40min after CCCP treatment are shown. Yellow outlined arrows point to colocalization between CARP2 (pseudo green color) and Rab7A or Rab5B (pseudo blue color) without mitochondria. Filled white arrows point to colocalization between CARP2 (pseudo green color), Mitochondria (pseudo red color) and Rab7A or Rab5B (pseudo blue color). Filled yellow arrows point to colocalization between CARP2 (pseudo green color) and Mitochondria (pseudo red color) without Rab5B. Scale 2μm, inset scale 1μm. (**C** and **D**) The graph represents the percentage of cells showing association of mitochondria with Rab7A (**C**) or Rab5B (**D**) and CARP2 (cells having more than four CARP2 and Rab7A/5B localized on MitoTracker were considered for counting). Error bars represent mean ± SEM in cases of (**C**) and (**D)** from three independent experiments with a minimum of 50 cells per experiment. Statistical significance was calculated using a two-tailed unpaired t-test. P-values are <0.0001 in the case of **C** and **D**.

**Figure 3.**
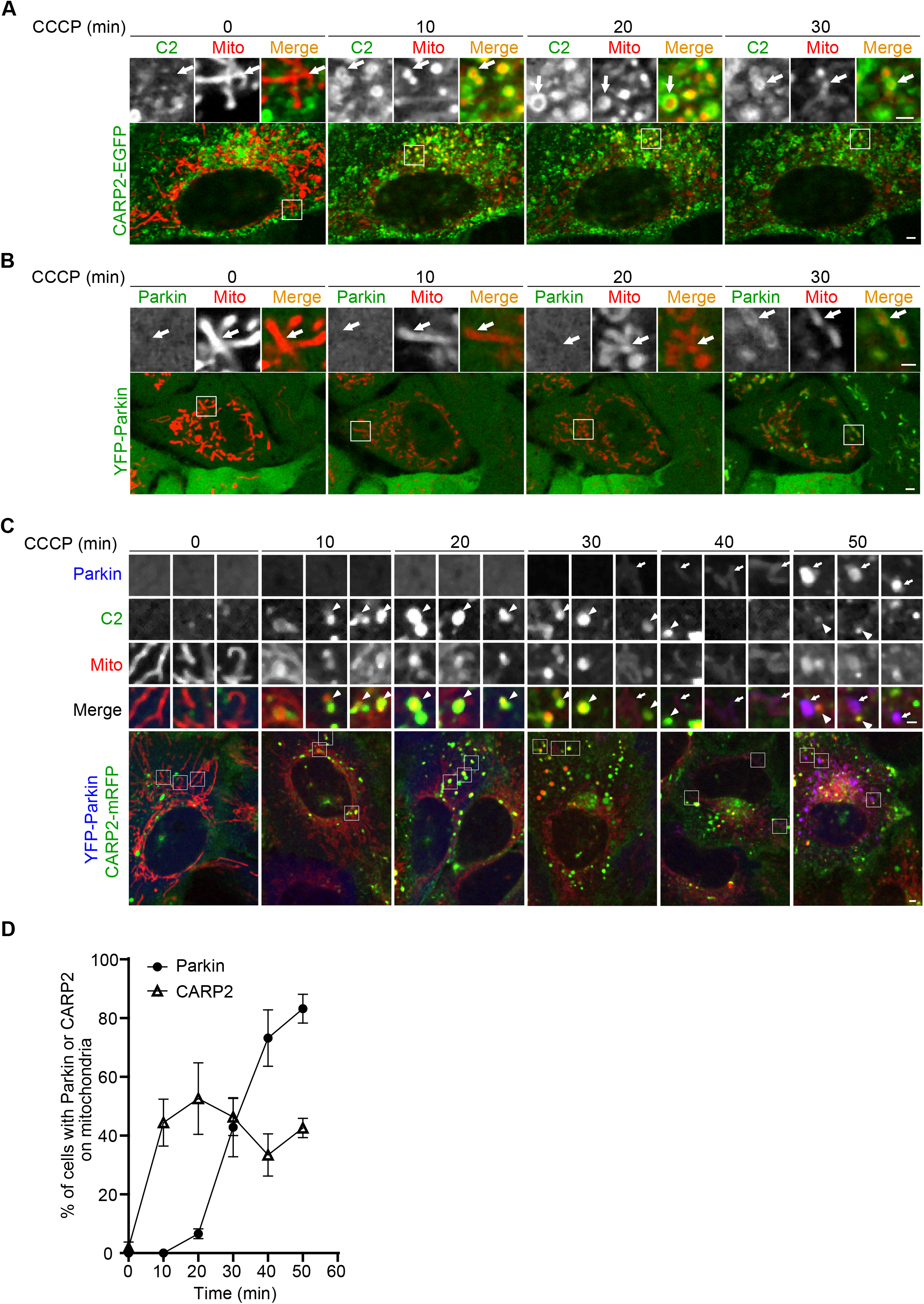
CARP2 associates with damaged mitochondria before Parkin. (**A** and **B**) A549 cells stably expressing CARP2-EGFP (**A**) or YFP-Parkin (**B**) stained with MitoTracker Red CMXRos for 30min were washed in DMEM followed by treatment with 20μM CCCP. Images of live cells at the indicated time points after treatment are shown Scale 2μm, inset scale 1μm. (**C**) U2OS cells stably expressing CARP2-mRFP and YFP-Parkin were pre-stained with MitoTracker Deep Red FM for 30min and washed in DMEM. These cells were subsequently treated with 20μM CCCP. Images of live cells at the indicated time points after treatment are shown. Colocalization between Parkin (pseudo blue) and Mitochondria (pseudo red) was shown with arrows. Colocalization between CARP2 (pseudo green) and Mitochondria (pseudo red) are shown with arrowheads. Scale 2μm, inset scale 1μm. (**D**) The graph represents the percentage of cells with mitochondria localized with either CARP2-EGFP or YFP-Parkin (cells having more than four CARP2 or Parkin localized on MitoTracker were considered for counting) at indicated time points. Data shown is from three independent experiments and a minimum of 50 cells were counted per experiment. Error bars represent mean ± SEM.

### CARP2-associated mitochondria are positive for Rab7A and Rab5B

Studies have shown that some of the endosomes or endosomal proteins are involved in the regulation of Parkin mediated mitochondrial clearance [6, 13, 15, 23]. Hence, we investigated whether CARP2-positive vesicles are associated with mitochondria, using some of the known mitophagy-linked endosomal markers, namely Rab5 and Rab7. Both Rab5 and Rab7 are widely used as markers to identify endosomes, and CARP2 is known to associate with intracellular vesicles that are positive for these endosomal markers [18, 19]. Towards this end, we have transiently transfected CARP2-mRFP stably expressing N2a cells with GFP-Rab5B or GFP-Rab7A and stained the mitochondria with Mitotracker Deep Red FM. Without any treatment, very little colocalization was noted between CARP2-positive Rab7A or Rab5B vesicles and mitochondria. However, upon CCCP treatment, a significant number of cells showed an association between CARP2-positive mitochondria and Rab7A, and to a lesser extent to Rab5B (Figure 2A, B). Quantification of the percentage of cells with CARP2 and Rab7A or Rab5B localized to mitochondria are shown in Figure 2C, D. These results demonstrate that CARP2-Rab7A/5B endosomes are associated with damaged mitochondria.

We further examined the dynamics of the association between endosomes and damaged mitochondria using time-lapse live imaging microscopy over a period of 12 min. Monitoring of cells stably expressing CARP2-EGFP and stained with MitoTracker Red CMXRos revealed the formation of protrusions of GFP-positive structures from CARP2 vesicles towards damaged mitochondria (Figure S1D and Movie 2), which transiently hover around mitochondria. To further understand this phenomenon, we have used a photobleaching strategy. A549 cells stably expressing CARP2-EGFP were exposed to bleaching using 488 nm Argon laser at a region of interest, consisting of CARP2-positive mitochondria. After bleaching, the movement of CARP2 vesicles (green) in the vicinity of damaged mitochondria (red) was monitored for a period of 2 minutes. As early as 41 seconds after photobleaching, movement of GFP-positive structures from the surrounding area of the bleached region towards the damaged mitochondria was noted (Figure S1E, Movie 3). As pointed by arrows, these structures appeared tubular in nature. Moreover, under these conditions, within two minutes the mitochondria were noted to be surrounded with multiple CARP2-positive vesicles, suggesting a plausible role for CARP2-positive endosomes in mitophagy. As these findings suggest that the CARP2 endosomes are associated with mitochondria, the role of CARP2 association with damaged mitochondria and Parkin, a well-known E3 involved in the elimination of damaged mitochondria, is explored further.

### CARP2 associates with damaged mitochondria prior to Parkin, and CARP2 KO cells show defective Parkin recruitment

Because mitophagy is known to be regulated by Parkin, we investigated the function of CARP2 in Parkin-mediated mitophagy. We monitored the localization of CARP2 and Parkin in CCCP treated A549 cells, stably expressing CARP2-EGFP or YFP-Parkin individually and pre-stained with MitoTracker Red CMXRos. As shown in Figure 3A, CARP2 localization to mitochondria was observed within 10min of treatment, whereas that of Parkin was noted only after 30min (Figure 3B). To investigate whether the kinetics of recruitment of both these proteins to mitochondria is specific to A549 cells, or similar pattern can be observed in other cells, we stably expressed both CARP2-mRFP and YFP-Parkin simultaneously in U2OS cell line. Simultaneous expression of these proteins also helped to monitor their relative recruitment vs mitochondria. As seen in A549 cells, CARP2 recruitment to mitochondria was found in early time points compared to Parkin recruitment. Surprisingly, very little colocalization between mitochondria that associates with CARP2 and Parkin were noted (Figure 3C). Time-course measurements indicated maximum localization of CARP2 within 10min, whereas Parkin peaked after 30min (Figure 3D). However, proximity between CARP2-(arrowheads) and Parkin-positive (arrows) mitochondria were observed, without overt localization (Figure 3C, 50min CCCP).

As CARP2 is localized to the mitochondria prior to Parkin, we were interested in exploring CARP2 function in regulating Parkin recruitment to the mitochondria. To investigate such an idea, we generated CARP2 knockout cells using the CRISPR-Cas9 system and used three independent colonies (# 2, 9 and 12) in our experiments. All three clones were validated using western blotting (Figure S2B). We monitored the status of Parkin localization to damaged mitochondria (stained with Mitotracker CMXRos) in CARP2 WT or KO cells stably expressing YFP-Parkin. In DMSO-treated cells, as expected, YFP-Parkin remained cytosolic in both CARP2 WT and KO cells (Figure S3, 4). In CCCP-treated WT cells, YFP-Parkin appeared as punctate structures that are positive for the MitoTracker, indicating its localization to damaged mitochondria. In CARP2 KO cells, however, a significant reduction in YFP-Parkin localized to mitochondria was noted with most of the Parkin remained diffused in the cytosol (Figure 4C, D), though the total amount of mitochondria in KO cells appeared similar to that found in WT. While differences in Parkin localization was significant up to 1h and at 6h of CCCP treatment, the differences were less pronounced during 2 to 4h suggesting a delay both in the recruitment and stabilization of Parkin on the mitochondria (Figure S3A, B) in KO cells. These observations from CARP2 KO cells (clone # 2) are not a result of the off-target effect, as we could observe this phenotype in two other independent CARP2 KO clones # 9 and 12 (Figure S2A). For all other experiments, CARP2 KO clone # 2 was used. These data suggest a plausible role of CARP2 in Parkin recruitment to mitochondria.

**Figure 4.**
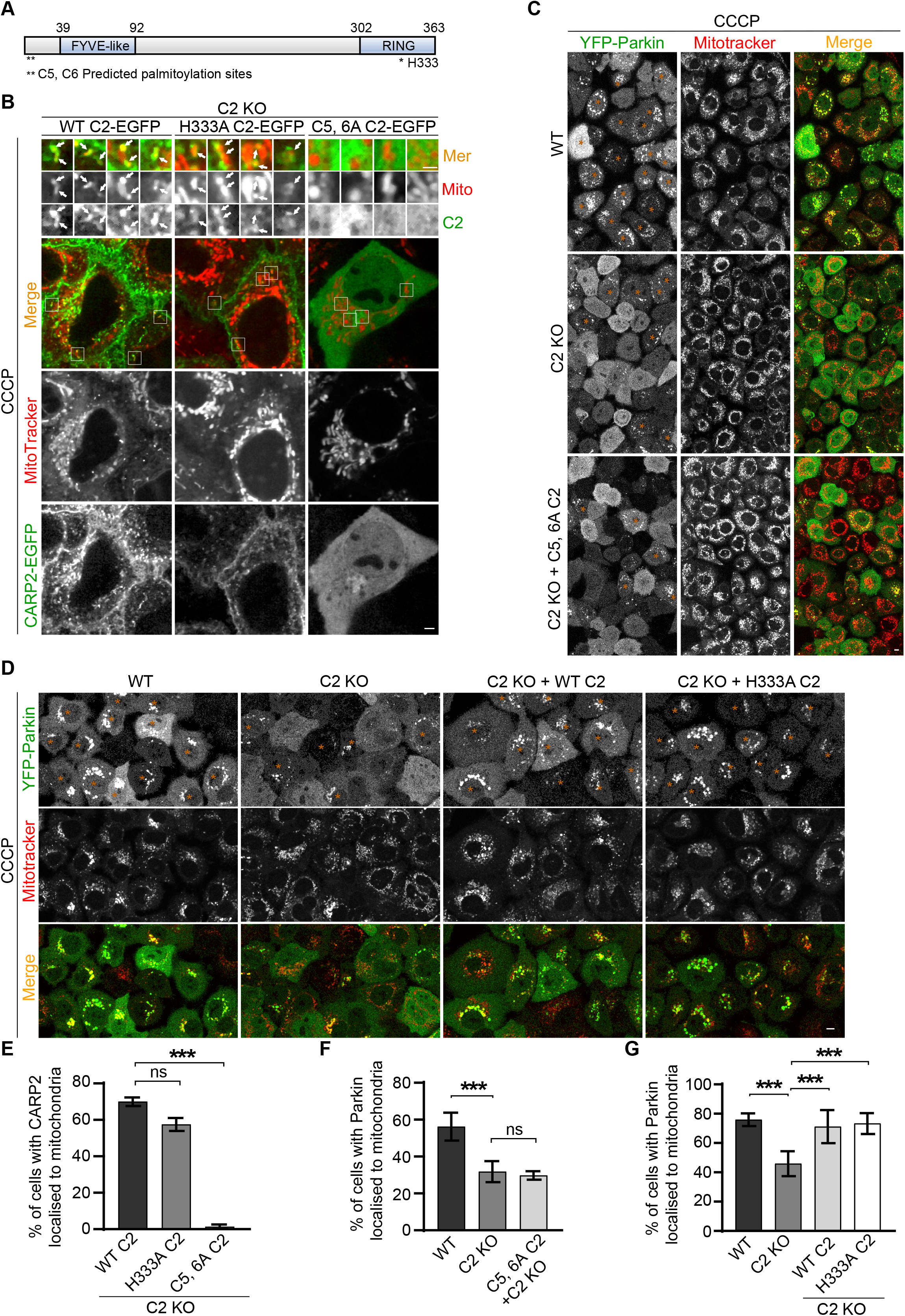
Endosomal-association-defective CARP2 variant fails to associate with mitochondria and affects Parkin recruitment. (**A**) Schematic representation of CARP2 with predicted palmitoylation sites (*cysteines at 5^th^ and 6^th^ positions) and a conserved histidine residue at 333^rd^ position in the RING domain. (**B**) A549 CARP2 KO cells expressing EGFP tagged CARP2 WT, or H333A, or C5, 6A mutant, were pre-stained with MitoTracker Red CMXRos. These cells were washed with DMEM and subsequently treated with 20μM CCCP. Representative images of live cells over 10-40min of treatment are shown. White arrows show the localization of CARP2 with mitochondria. Inset contrast was adjusted for better visibility. Scale 2μm, inset scale 1μm. (**C, D**) YFP-Parkin stably expressing A549 WT, or CARP2 KO, or CARP2 KO reconstituted with untagged CARP2 WT, or C5, 6A, or H333A were pre-stained with MitoTracker Red CMXRos for 30min. These cells were washed with DMEM and subsequently treated with 10μM CCCP. Representative images of live cells at 6h (5.5h to 6.5h) of treatment are shown. Orange asterisks represent cells counted as positive for Parkin on mitochondria. Scale 5μm. (**E**) The bar graph shows the percentage of cells with more than four CARP2-EGFP puncta localized to MitoTracker. Error bars represent mean ± SEM from three independent experiments. A minimum of 50 cells were counted for each experiment. Statistical significance was determined using ordinary one-way ANOVA followed by Tukey’s multiple comparisons test. P-values for WT C2 vs. H333A C2 is 0.1063 and for WT C2 vs. C5, 6A C2 is <0.0001. (**F, G**) Bar graph showing the percentage of cells with at least fifteen YFP-Parkin puncta localized to MitoTracker during this period. Data shown are from three independent experiments and a minimum of 75 cells were counted per experiment. Error bars represent mean ± SEM. Statistical significance was calculated using ordinary one-way ANOVA followed by Tukey’s multiple comparisons test. P-values for C2 KO vs WT is 0.0001 (**F**), C2 KO vs C2 KO + C5, 6A is 0.9321, C2 KO vs WT is < 0.0001 (**G**), C2 KO vs C2 KO + WT C2 is 0.0006, and C2 KO vs. C2 KO + H333A C2 is 0.0003.

### Endosomal-association-defective CARP2 variant fails to associate with mitochondria and affects Parkin recruitment

CARP2, in addition to the ubiquitin ligase domain, consists of two cysteine residues (C5, 6) that are predicted to be involved in palmitoylation (Figure 4A). Both these cysteines are required for the endosomal association as CARP2 C5, 6 A variant is unable to localize to intracellular vesicles (Figure 4B and [19, 24]). To understand whether ubiquitin ligase activity and/or endosomal localization are required for the association of CARP2 with damaged mitochondria, we have reconstituted A549 CARP2 KO cells with EGFP tagged CARP2 variants. The reconstituted cells were stained with MitoTracker Red CMXRos, and the association of various CARP2 mutant variants with damaged mitochondria was assessed after CCCP treatment. Interestingly, CARP2 ubiquitin ligase inactive mutant (H333A) also showed colocalization with damaged mitochondria after CCCP treatment. The H333A mutant association with mitochondria appears to be similar to that noted with WT CARP2 with some differences. CARP2 WT formed a distinct ring-like pattern around the damaged mitochondria, which was not apparent in the case of the H333A mutant (Figure 4B). These results suggest that ubiquitin ligase activity is not essential for CARP2 association with mitochondria.

Unlike WT and H333A that localized to intracellular vesicles, endosomal localization defective CARP2 variant (C5, 6A) showed a diffused cytosolic pattern. Moreover, in CCCP treated cells, the C5, 6A variant showed very little detectable association with damaged mitochondria (Figure 4B). Quantification of the percentage of cells with CARP2 variants on mitochondria is included in Figure 4E. These observations suggest predicted palmitoylation residues are essential for CARP2 association with damaged mitochondria, as a change of these conserved cysteines affected not only endosomal localization of CARP2 but also its association with mitochondria.

Next, to understand whether the association of CARP2 with mitochondria is required for Parkin recruitment, we have generated CARP2 KO cells reconstituted with untagged versions of CARP2 WT, or H333A or C5, 6A along with YFP-Parkin. These cells were stained with MitoTracker CMXRos, and Parkin localization was observed in DMSO- and CCCP-treated conditions. As expected, Parkin appeared as a diffused cytosolic pattern with no association with MitoTracker in all the cells after DMSO treatment (Figure S4). Whereas KO cells expressing either WT or H333A showed an increase of Parkin recruitment compared to the KO cells, but similar to A549 cells (Figure 4D, G), KO cells reconstituted with C5, 6A mutant showed a pattern similar to that noted in KO alone cells (Figure 4C, F). These results suggest the importance of CARP2 association with mitochondria for Parkin recruitment. Interestingly, these results also point out that CARP2 facilitates Parkin recruitment to the damaged mitochondria independent of its ligase activity. However, CARP2 requires N-ter cysteine residues, which are important for recruiting Parkin to damaged mitochondria.

### CARP2 interacts with Parkin and regulates mitophagy independent of its ubiquitin ligase activity

Since the data indicated that CARP2 facilitates Parkin recruitment to the mitochondria, we investigated whether CARP2 can interact with Parkin. To explore such an idea, we have transfected HEK293T cells with cDNA constructs of Parkin and CARP2 WT or H333A mutant with appropriate controls. The cellular lysates were used to immunoprecipitate CARP2 using CARP2 specific antibody and the immunoprecipitates were probed for Parkin. In comparison to the control samples, we were able to consistently see the presence of Parkin in the precipitates from CARP2 WT samples. Surprisingly, the amount of Parkin pulled was much more in the precipitates from the lysates of cells overexpressing ligase inactive mutant than of WT CARP2 though the amount of CARP2 present in both WT and mutant samples were similar (Figure 5A). As these results suggested that CARP2 can interact with Parkin, to further validate the same, we have performed the immunoprecipitation of endogenous CARP2 in A549 cells stably expressing YFP-Parkin. Consistent with the data from overexpression, in untreated cells, very little basal level of interaction between endogenous CARP2 and Parkin was noted. Moreover, after CCCP treatment, a noticeably more amount of Parkin was observed in the immunoprecipitates of CARP2, suggesting an increased association between these proteins in treated cells (Figure 5B). This interaction is specific, as no Parkin band was detected in immunoprecipitates from the lysates of CCCP-treated CARP2 KO cells (Figure 5B).

**Figure 5.**
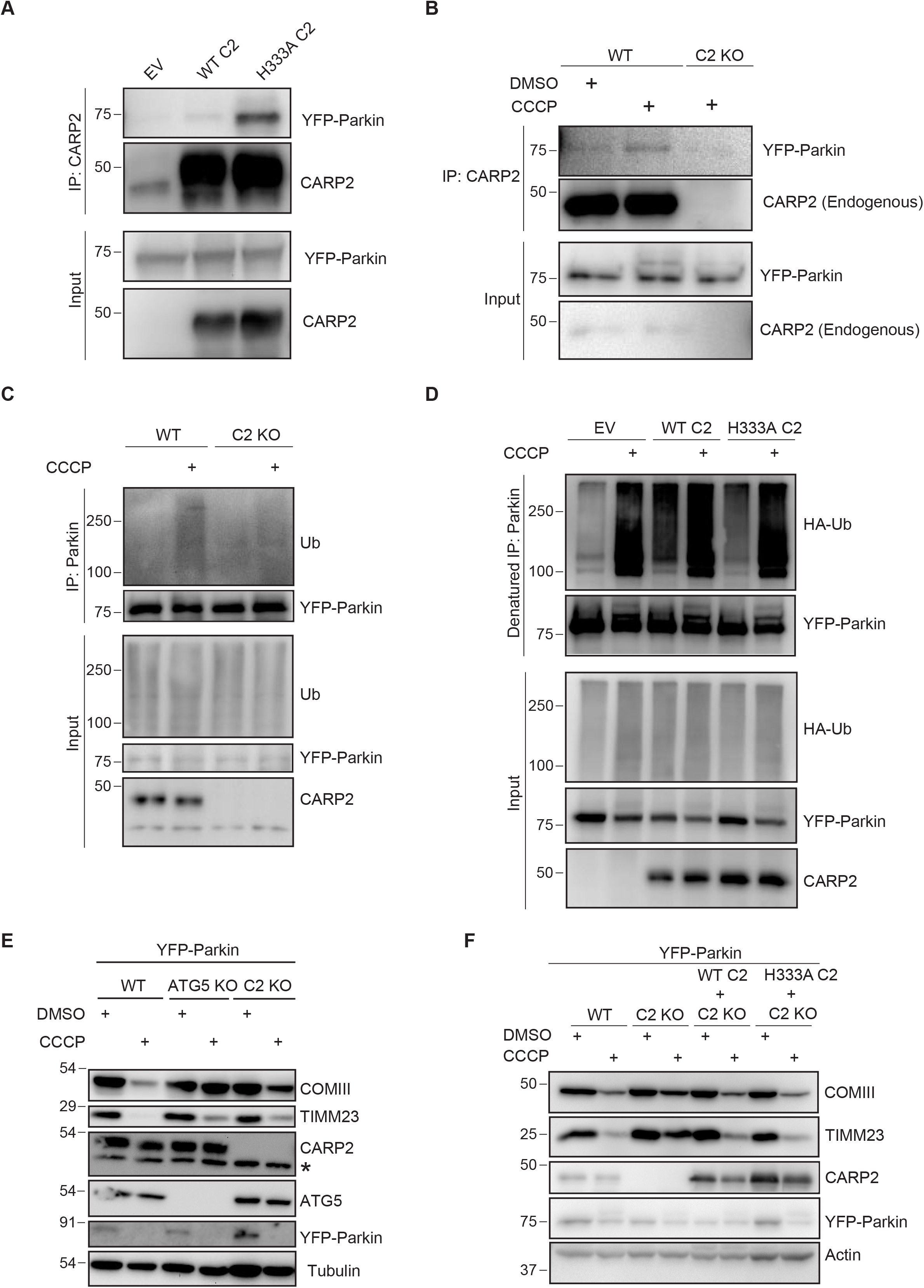
CARP2 interacts with Parkin and regulates mitophagy independent of its ubiquitin ligase activity. (**A**) HEK 293T cells transfected with empty vector (EV) or Flag-WT CARP2 or Flag-H333A CARP2 were transiently cotransfected with YFP-Parkin. Cells were lysed and CARP2 was immunoprecipitated using CARP2 specific antibody. The precipitates and lysates were probed with indicated antibodies. (**B**) A549 WT or CARP2 KO cells stably expressing YFP-Parkin were treated with DMSO or 10μM CCCP for 3h. Cells were lysed and endogenous CARP2 was immunoprecipitated using CARP2 specific antibody. The precipitates and lysates were probed with indicated antibodies. (**C**) A549 Wild Type (WT) or CARP2 KO cells stably expressing YFP-Parkin were treated with DMSO or 10μM CCCP for 3h. Cells were lysed and YFP-Parkin was immunoprecipitated using Parkin specific antibody. The precipitates and lysates were probed with indicated antibodies. (**D**) HEK 293T cells transiently transfected with empty vector (EV) or CARP2 WT or H333A along with HA-Ub and YFP-Parkin were treated with DMSO or 10μM CCCP for 3h. Cells were lysed and YFP-Parkin was immunoprecipitated using a Parkin specific antibody under denaturation conditions. The precipitates and lysates were probed with indicated antibodies. (**E**) A549 cells (WT or ATG5 KO or CARP2 KO) stably expressing YFP-Parkin were treated with DMSO or 20μM of CCCP for 24h. The cell lysates were subjected to western blotting with the indicated antibodies. * denotes a nonspecific band. (**F**) A549 cells WT, or CARP2 KO, or CARP2 KO reconstituted with untagged CARP2 WT or H333A stably expressing YFP-Parkin were treated with DMSO or 10μM of CCCP for 24h. The cell lysates were subjected to western blotting with the indicated antibodies.

As CARP2 can interact with Parkin and facilitates Parkin recruitment to the mitochondria, we were interested to know whether CARP2 promotes Parkin ubiquitination. For this, we have accessed the extent of ubiquitination in Parkin precipitates from lysates of A549 WT and CARP2 KO cells stably expressing YFP-Parkin with or without CCCP treatment. Parkin was immunoprecipitated from the cell lysates using Parkin specific antibody, and the precipitates were probed separately with Parkin antibody and anti-Ub antibody. The amount of Parkin that was precipitated remained more or less the same in all the samples. Under DMSO treated conditions, the amount of polyubiquitination on Parkin pull-down from both WT and KO cells remained the same. As expected, an increase in polyubiquitination was observed upon CCCP treatment in WT cells, whereas this increase was substantially reduced in cell lysates that do not express CARP2 (Figure 5C), suggesting that CARP2 contributes to the ubiquitination of Parkin under CCCP-treated conditions. To investigate whether the increased polyUb signal in Parkin immunoprecipitate is indeed on Parkin, we have transfected HEK293T cells with cDNAs of CARP2 variants or Parkin along with HA-Ub. The extent of polyubiquitination on Parkin was assessed by performing immunoprecipitation under denaturation conditions using Parkin specific antibody. In DMSO treated conditions, Parkin was found to be substantially more polyubiquitinated in the presence of WT CARP2 compared to the empty vector control and the ligase inactive mutant. Notwithstanding a substantial increase in the extent of polyubiquitinated Parkin upon CCCP treatment from control cells, a noticeable enhancement in polyubiquitination was observed in CARP2 WT expressing cells (Figure 5D). These results collectively suggest that CARP2 can interact with Parkin and promotes its polyubiquitination.

As we found that CARP2 vesicles surrounds damaged mitochondria and influences Parkin recruitment to mitochondria, we investigated the role of CARP2 in the elimination of mitochondria. For this, we monitored levels of two mitochondrial proteins, Complex III core I (COMIII), IMM/matrix protein or TIMM23, an inner mitochondrial membrane (IMM) protein [25, 26] with or without CCCP treatment. Consistent with the report of elimination of the mitochondria upon CCCP treatment [25], a significant decrease in the levels of COMIII and TIMM23 was noted in A549 cells. Importantly, the extent of reduction of both these proteins were much less in the absence of CARP2, i.e., in CARP2 KO cells after 24 h of CCCP treatment, suggesting that CARP2 contributes to the elimination of damaged mitochondria (Figure 5E). This reduction was similar to that observed in ATG5 KO cells (Figure 5E), where Parkin-dependent mitophagy is known to be ineffective [9]. To further validate the role of CARP2 in Parkin mediated clearance of damaged mitochondria, we have monitored the levels of COMIII or TIMM23 in A549 CARP2 KO cells reconstituted with WT or ligase inactive mutant. Concomitant with the normal Parkin recruitment to mitochondria observed in these cells (Figure 4D), expression of both WT and ligase inactive mutant rescued the degradation of mitochondrial proteins, TIMM23 and COMIII (Figure 5F). These results collectively indicate the crosstalk between endosomal ubiquitin ligase CARP2 and cytosolic ubiquitin ligase Parkin in the clearance of damaged mitochondria.

## Discussion

Endosomes are believed to provide membranes for the autophagosomes and regulate autophagy [27]. Emerging evidence envisages important roles for endosomal-associated proteins in the mitochondrial clearance by lysosomes via ESCRT pathway or Parkin-mediated mitophagy [6, 13, 23]. Post-translational modification with ubiquitin (ubiquitination) mediated by ubiquitin ligases like Parkin plays a critical role in maintaining mitochondrial homeostasis. Parkin association with mitochondria while involves PINK1, and the PINK1/Parkin feedforward loop-generated phospho-Ub facilitates activation and multimerization/stabilization of Parkin on mitochondria [10, 28, 29], many details of the role of endosomes in this process is not known. Our study proposes a novel mechanism on how endosomes/endosomal associated molecules regulate Parkin recruitment and elimination of damaged mitochondria. In this study, we demonstrated that CARP2-associated endosomes surround damaged mitochondria. We provide evidence that a variant of CARP2, C5, 6A mutant - that is deficient in its endosomal localization is unable to associate with mitochondria. Moreover, we found that mitochondria-associated with CARP2 vesicles are also positive for Rab7A, or to a lesser extent Rab5B, long used as endosomal markers. Collectively, these results indicate that it is the CARP2 endosomal vesicles, not CARP2 molecules that associate with damaged mitochondria. The mechanism(s) by which CARP2-endosomes sense damaged mitochondria and move towards organelle needs to be explored.

Another exciting aspect of CARP2 association with mitochondria is that it was observed as early as 10min of CCCP treatment, prior to Parkin translocation to mitochondria. Moreover, both CARP2- and Parkin-positive mitochondria appeared as distinct populations with very few mitochondria exhibiting association with both CARP2 and Parkin at the same time. However, it was also noted that several of the CARP2- and Parkin-positive vesicles are in close proximity. The existence of distinct mitochondrial populations based on membrane potential and Parkin translocation was reported earlier [9]. We believe that these discrete populations represent different, but yet connected stages, of organelle disintegration that are reported during mitophagy [7, 30, 31]. These results, along with the observation of compromised Parkin recruitment in CARP2 KO cells, led us to propose that the association of CARP2 vesicles with the mitochondria prime these organelles for Parkin recruitment and this process involves separation of CARP2 vesicles with some mitochondrial content.

We also provide evidence that CARP2 can interact with and facilitate the ubiquitination of Parkin in CCCP-treated cells. Interestingly, the ligase activity of CARP2 is not required for interaction with Parkin or the recruitment of later to and elimination of mitochondria. Such an E3-independent role in autophagy is not limited to CARP2 alone. Other ubiquitin ligases, namely TRIM13 and Smurf1, that regulate autophagy are reported to function independently of their respective ubiquitin ligase activities [32, 33]. Smurf1 in addition to HECT’s domain that confers ubiquitin ligase activity also possesses a phospholipid-binding C2 domain that anchors to membrane phospholipids and functions in protein targeting to the subcellular compartments [34]. While Smurf1 regulates mitophagy independent of its E3 ligase activity, its C2 domain is required for the same. Similar to Smurf1, CARP2 is an E3 with an FYVE-like domain, that is believed to bind to phospholipid molecules with the help of post-translational modifications such as palmitoylation [16, 24]. CARP2 has been shown to be palmitoylated [24], and the sites for this modification are predicted to be Cysteine residues in positions 5 and 6. CARP2 (C5, 6A) mutant not only failed to associate with damaged mitochondria but also was unable to rescue Parkin translocation to the mitochondria in CARP2 KO cells. Though both CARP2 and Smurf1 function independent of their E3 ligase activity, they require domains that are responsible for lipid binding to regulate mitophagy, suggesting both these proteins might work in a similar fashion in regulating mitophagy. As recycling endocytic vesicles are shown to contribute to autophagosomes formation and mitophagy [15], further studies in this area further our understanding of the biogenesis of autophagosomes in Parkin-mediated mitophagy. An important role of CARP2 in mitophagy is further supported by reports of modulation of Parkin recruitment to mitochondria and its activity by two other molecules p53 and MDM2 [35, 36], though many details of this process remain unknown. Importantly, both p53 and MDM2 are interacting partners of CARP2 [37]. Further investigations are required to exactly understand roles played by CARP2 vs MDM2/p53 in mitophagy.

The proposed model in Figure S5 is based on observations from our study and the literature. This study also opens many important aspects for further studies. How does CARP2-endosomes sense damaged mitochondria and the role of PINK1 in CARP2 recruitment to damaged mitochondria? Another important issue that warrants further exploration is the role of CARP2 in Parkin independent mitophagy and whether CARP2-positive mitochondrial vesicles are linked to events in the scheme of piecemeal mitophagy that is believed to occur at the physiological level. More studies are needed to understand the nature and function of CARP2-positive mitochondrial vesicles. This study with CARP2 would further unravel the contribution of endosomes to Parkin-dependent mitophagy. The role for CARP2-positive endosomes may not be limited to mitophagy, but might also extend to various other fields autophagy like aggrephagy, pexophagy, reticulophagy, xenophagy, nucleophagy and zymophagy.

## Materials and methods

### Cell culture and generation of stable cell lines

All cells were maintained in Dulbecco’s modified Eagle’s medium (DMEM plus Glutamax Gibco Cat # 10569-010) containing 10% heat-inactivated FBS (Gibco Cat # 10270-106) and 1% penicillin-streptomycin (HiMedia Cat # A001A) in a 5% CO2 incubator at 37°C. For inducing mitochondrial damage, 80-85% of confluent cells were treated with Carbonyl cyanide-m-chlorophenylhydrazone (CCCP) (Sigma, Cat # C-2759) or Valinomycin (Sigma, Cat # V0627) after 18h of plating. Mitochondria were stained with 100nM of either MitoTracker Red CMXRos (Invitrogen, Cat # M7512) or MitoTracker Deep Red FM (Invitrogen, Cat # M22426) for 30min in culture media followed by one wash in DMEM before CCCP treatment. To develop U2OS cells stably expressing YFP-Parkin and EBFP2-Mito-7, initially, the cells were transfected with YFP-Parkin and stable cells were generated by selection using G418 (Sigma, Cat # A1720). The stably expressing cells were further transfected with EBFP2-Mito-7 and cells expressing both the transgene were sorted using flow cytometer sorter equipped with a 375-nm laser for EBFP excitation (FACS Aria III). Multiple clones were expanded after further selection in G418 for two weeks, and stably expressing clones were used for the current study. Cells stably expressing CARP2-EGFP were generated as per the previous protocol [19]. Briefly, HEK293T cells were transfected with pMYsIP-CARP2-EGFP along with pCAG-VSVG and pUMVC plasmids using Lipofectamine 3000 reagent (Invitrogen Cat # L3000-015). Four days post-transfection, cell supernatants were harvested and were incubated with A549 or U2OS cells in the presence of 4μg/mL Polybrene (Sigma, Cat # H9268). Cells were cultured in Puromycin (1μg/mL) (Sigma Cat # P8833) containing DMEM for selection. Other stable cell lines mentioned were also prepared using a similar protocol.

### Plasmids, Antibodies and Reagents

Human CARP2 cDNA (NCBI reference sequence: NM_001017368.2:141-1232) or YFP-Parkin [9] with indicated mutations and tags were cloned to pMYs-IP for stable cell line generation [20]. pFLAG-CMV2 CARP2 WT and H333A was a gift from Wafik S. El-Deiry [16]. Following constructs used were received from Addgene, YFP-Parkin was a gift from Richard Youle (Addgene plasmid # 23955) [9], GFP-Rab5B and GFP-Rab7A were a gift from Gia Voeltz [38], EBFP2-Mito-7 was a gift from Michael Davidson (Addgene plasmid # 55248) [39], pCAG-VSVG was a gift from Arthur Nienhuis and Patrick Salmon (Addgene plasmid # 35616), pUMVC was a gift from Bob Weinberg (Addgene plasmid # 8449) [40], PX458 was a gift from Feng Zhang (Addgene plasmid # 48138) [41]. Following antibodies were used in this study, RFP antibody (Chromotek Cat # 5f8-100), Polyclonal CARP2 antibody raised in Rabbit (Generated in our laboratory), GAPDH (Abgenex Cat # 10-10011), Beta Actin (Cat # E-AB-20058), Tubulin (Sigma-Aldrich Cat # T6199), ATG5 (CST Cat # 12994T), UQCRC1 (For Complex III, Core I, Invitrogen Cat # 16D10AD9AH5), TIMM23 (SCBT Cat # sc-514463), GFP (Mouse Living Colours Clonetek Cat # 632375), GFP (Invitrogen Cat # A6455)Parkin (SCBT Cat # sc-32282), Anti-mouse HRP (Invitrogen Cat # 61-6520), Anti-rabbit HRP (Invitrogen Cat # 65-6120), Anti-rat HRP (Jackson ImmunoResearch Cat # 712-035-150).

### Western Blotting

18h after plating, cells at 80-85% confluency were washed and pelleted at 850×g in ice-cold PBS. Cell lysis was done in ice for 30min using lysis buffer containing 50mM Tris-Cl pH 7.5, 150mM Sodium chloride, 1% TritonX-100, 1mM EDTA pH 8, 1mM Phenylmethylsulfonyl fluoride (PMSF Cat # P7626) and 1X Protease inhibitor cocktail (Sigma Cat # P8340). The supernatant was collected after centrifuging the lysate at 13500×g for 10min at 4°C. Total protein levels in the supernatant were normalized using BCA (Pierce BCA Protein Assay Kit Cat # 23225). Samples were mixed with Laemmli buffer and heated for 10min at 95°C before loading them into SDS-PAGE. Samples separated in SDS-PAGE were later transferred to the PVDF membrane (Merck Cat # IPVH00010). Immunoblot analysis was performed by using antibodies as indicated. They were visualized with chemiluminescent HRP substrate (Merck Cat # WBKLS0500). Signal intensities were calculated using Quantity One version 4.6.9.

### Immunoprecipitation

Cell extracts were prepared in IP buffer containing 20mM Tris-Cl pH 7.6, 150mM Sodium Chloride, 5mM EGTA, 0.5% TritonX-100, 1mM Phenylmethylsulfonyl fluoride, 1X Protease inhibitor cocktail (Sigma Cat # P8340 and Cat # 04693159001), 1X Phosphatase inhibitor cocktail (Sigma Cat # P5726 and # P0044), and 100mM N-Ethylmaleimide (Sigma Cat # E3876). Lysates were then incubated with CARP2 antibody at 4°C for 6h. The CARP2 or Parkin protein complex was then pulled down by incubating antibody-containing lysate with Protein A or G agarose beads at 4°C for 1h. Agarose beads were suspended in Laemmli buffer and heated at 95°C in all samples or at 37°C in the case of Parkin IP samples. Immunoprecipitation under denaturation condition was performed by lysing the cells at 90°C for 10min in lysis buffer containing 10mM Tris-Cl pH 7.6, 1% SDS, 5mM EDTA, 10mM DTT, 1mM Phenylmethylsulfonyl fluoride, 1X Protease inhibitor cocktail, 1X Phosphatase inhibitor cocktail, and 100mM N-Ethylmaleimide. The lysate was then diluted 10 times with the IP buffer and Immunoprecipitation was performed as mentioned earlier.

### CRISPR KO preparation

A549 cells were transfected with PX458 containing CARP2 or ATG5 gRNAs using Lipofectamine 3000 and sorted for GFP-positive cells after 3 days of transfection using flow cytometry. Later, cells were serially diluted and plated in 96 wells to grow single colonies. Knockout of CARP2 or ATG5 from each clone was confirmed using western blot. Sequences of gRNA used are as follows: CARP2 5’-GGTGGAACCATGCTGGAGTG-3’ and ATG5 5’-AACTTGTTTCACGCTATATC-3’ [35].

### Live cell imaging and immunocytochemistry

Live cell imaging was performed after 18h of plating of cells in glass-bottom dishes (MatTek 254 Corporation, Cat # P35G-1.0-20-C). 14 to 18 h of CCCP were chosen for time-lapse, photobleaching experiments and super-resolution imaging because early hours of CCCP showed the dynamic movement of CARP2 vesicles which made the capture of CARP2-mitochondria population and photobleaching technically difficult. These movements were slower at later hours of CCCP. Glass bottom dishes were placed into an on-stage incubator with 5% CO_2_ and 37°C. For immunocytochemistry, cells were grown on coverslips and fixed in 4% Paraformaldehyde (PFA) for 14min. Permeabilization was done in 0.1% Triton X-100 for 10min followed by blocking in 3% BSA for 30min. Cells were then incubated with anti-GFP and anti-TOMM20 antibodies followed by incubating with respective secondary fluorophore-conjugated antibodies; Alexa Fluor 488 donkey anti-Rabbit (Invitrogen Cat # A-21206) and Alexa Fluor 568 goat anti-mouse (Invitrogen Cat # A-11031). After the antibody incubation, cells were mounted on a glass slide with ProLong Gold antifade reagent (Invitrogen, Cat # P36930). Images were acquired using Leica TCS SP5 II laser scanning inverted confocal microscope with oil-immersion objectives (HCX PL APO CS 63.0×1.40 OIL UV or HCX PL APO CS 100.0×1.40 OIL). Photobleaching was carried out using a 488 nm argon laser at 80% laser power on a chosen ROI (Region Of Interest). Representative confocal images used for figure preparation were noise reduced using LAS AF median filtering, keeping the same settings for all samples. Unprocessed images were used for all quantifications. Super-resolution imaging was carried out using Structured Illumination Microscopy (3D-SIM), N-SIM from Nikon (Tokyo, Japan). Cells grown in a glass-bottom dish were maintained at 5% CO2 with 37°C using an on-stage incubator from Tokai Hit (Japan). Cells were imaged using 100 x 1.49 NA oil immersion objective. Images were taken using EMCCD Camera iXon 897 (Andor, USA) were reconstructed using NIS elements software following standard protocol.

## Statistical analysis

Data were collected from at least three independent experiments and are shown as mean ± SEM as indicated. Significance levels were set at ∝ = 0.05. *P < 0.05, **P < 0.01, and ***P < 0.001. All statistical analyses were performed in GraphPad Prism (Version 8.3.1).

## Acknowledgements

We wish to thank DST-INSPIRE for the fellowship of RR, DBT for the fellowship of AKV and IISER TVM for the fellowship of NDN. This project is partly funded by Department of Biotechnology support for lifetime imaging facility awarded to TRS, Department of Biotechnology grant (BT/PR21325/BRB/10/1554/2016), Department of Science and Technology-Science and Engineering Research Board (DST-SERB) grant (EMR/2016/008048) and IISER TVM intramural funding awarded to SMS. We also thank Nayan Suryawanshi, Rithwik P. Nambiar, Abhishek Raghunathan and Dr. Ajitha T. K. for their technical help, Anurup KG for super-resolution microscopy imaging support, Surabhi S. V. and Tilak Prasad for their technical help in cell sorting by FACS, and Prof. M. K. Mathew for helpful discussion during manuscript preparation.

## Author contributions

RR, AKV, NDN and AC performed the experiments and acquired the data. RR, AKV, NDN, SMS and TRS critically analyzed the data. RR, AKV, NDN and SMS wrote the manuscript. RR, AKV, NDN and SMS conceived the project and planned the experiments.

## Abbreviations

CARP2: Caspase-8- and -10-associated RING Protein 2
CCCP: Carbonyl cyanide 3-chlorophenylhydrazone
EGFP: Enhanced Green Fluorescence Protein
KO: Knockout
UT: Untreated
WT: Wild-type

## Figure legends

**Supplemental figure S1.**
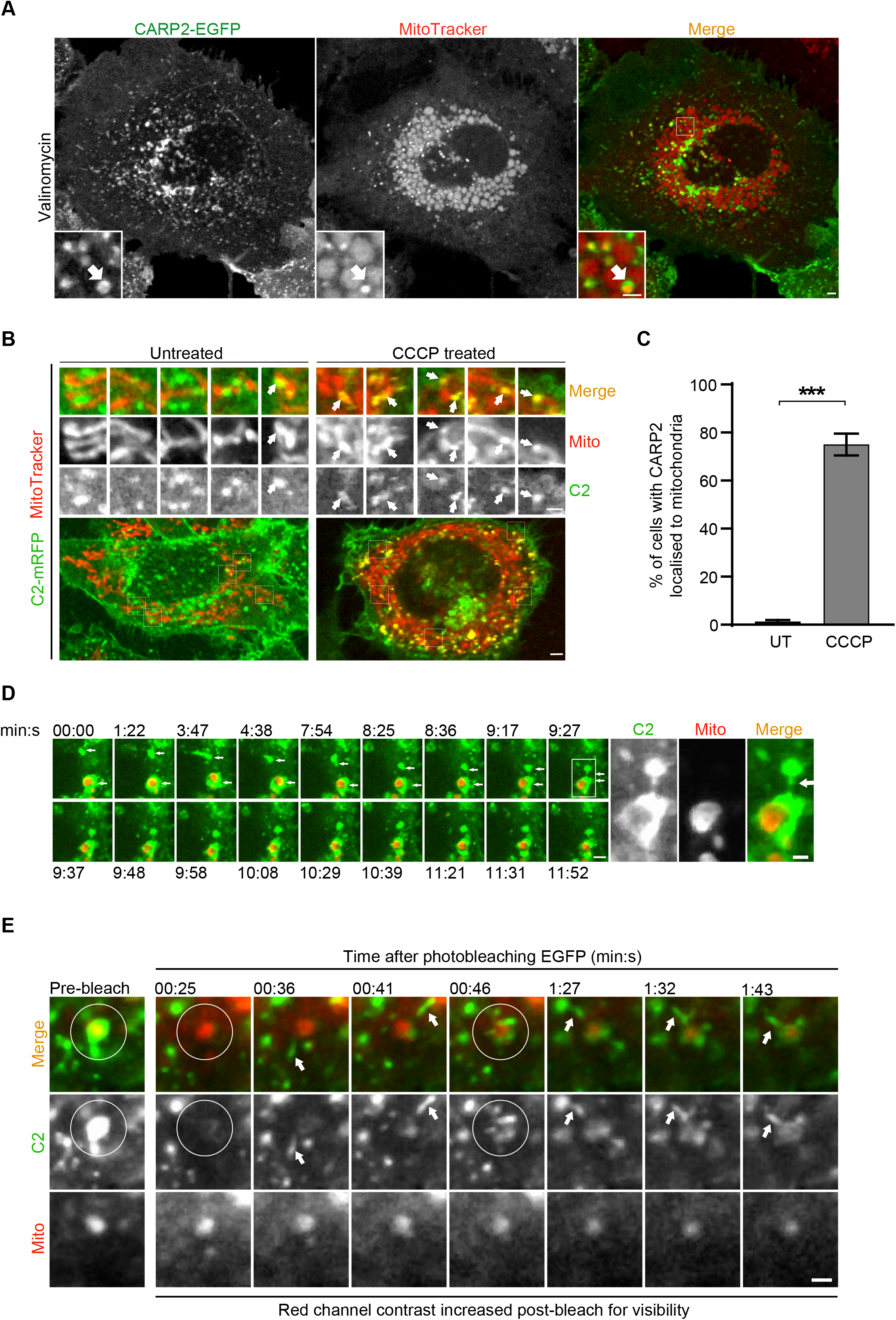
Dynamic association of CARP2-positive endosomes with damaged mitochondria. (**A**) A549 cells stably expressing CARP2-EGFP were pre-stained with MitoTracker Red CMXRos and subsequently treated with 20μM valinomycin. Images were acquired after 3.5h treatment. Scale 2μm, inset scale 1μm. (**B**) N2a cells stably expressing CARP2-mRFP were pre-stained with MitoTracker Red CMXRos and the cells were washed in DMEM and subsequently treated with 10μM CCCP. Images were acquired during 10min to 1h of treatment. Inset contrast was adjusted for better visibility. Scale 2μm, inset scale 1μm. (**C**) The bar graph shows the percentage of cells with more than four CARP2-EGFP puncta localized to MitoTracker. Error bars represent mean ± SEM from three independent experiments. A minimum of 50 cells were counted for each experiment. Statistical significance was calculated using a two-tailed unpaired t-test. p <0.0001. (**D**) A549 stably expressing CARP2-EGFP cells were stained for 30min with MitoTracker Red CMXRos and subsequently washed with DMEM. These cells were treated for 14h of CCCP (20μM) treatment (chosen frames). Time-lapse images of these cells were taken as indicated. White arrows indicate CARP2-EGFP-positive vesicles in proximity with MitoTracker Red CMXRos at the indicated time points, scale 2μm and inset scale 1μm. These results are also presented as a movie (**Movie 2**). (**E**) A549 stably expressing CARP2-EGFP cells were stained for 30min with MitoTracker Deep Red FM and subsequently washed with DMEM. These cells were treated with CCCP (20μM) for 18h. Cells with CARP2-EGFP around MitoTracker Deep Red FM were imaged live before and after photobleaching (chosen frames are shown). Circles show the region of photobleaching. Arrows show the movement of CARP2-EGFP-positive vesicles towards the photobleached region. The contrast of the MitoTracker panel alone is adjusted in frames after photobleaching for better visibility. CARP2-EGFP settings are kept the same before and after photobleaching. Scale 1μm. The same results are presented as a movie (**Movie 3**). Frames from 10 s to 20 s of the movie shows zoomed ROI during photobleaching, which is a software automated image processing.

**Supplemental figure S2.**
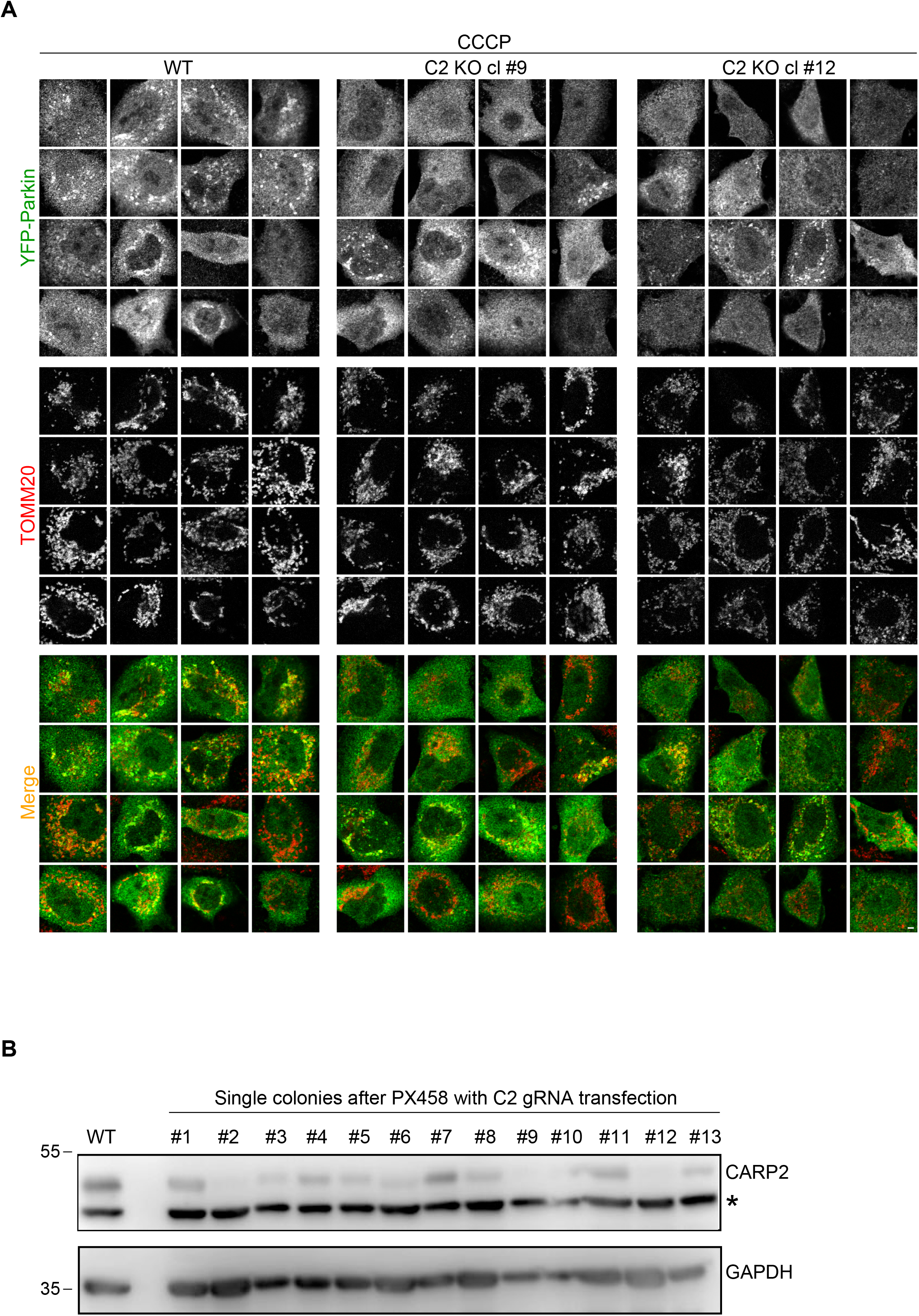
Impaired Parkin recruitment phenotype in two independent CARP2 KO clones. (**A**) Wild-type or CARP2 KO clones (# 9 or 12) A549 cells transiently transfected with YFP-Parkin were treated with 10μM CCCP. Images were captured after immunostaining the cells with anti-GFP and anti-TOMM20 antibodies. (**B**) Western blot showing the screening for CRISPR-Cas9 KO of CARP2 in A549 cells. * denotes a nonspecific band.

**Supplemental figure S3.**
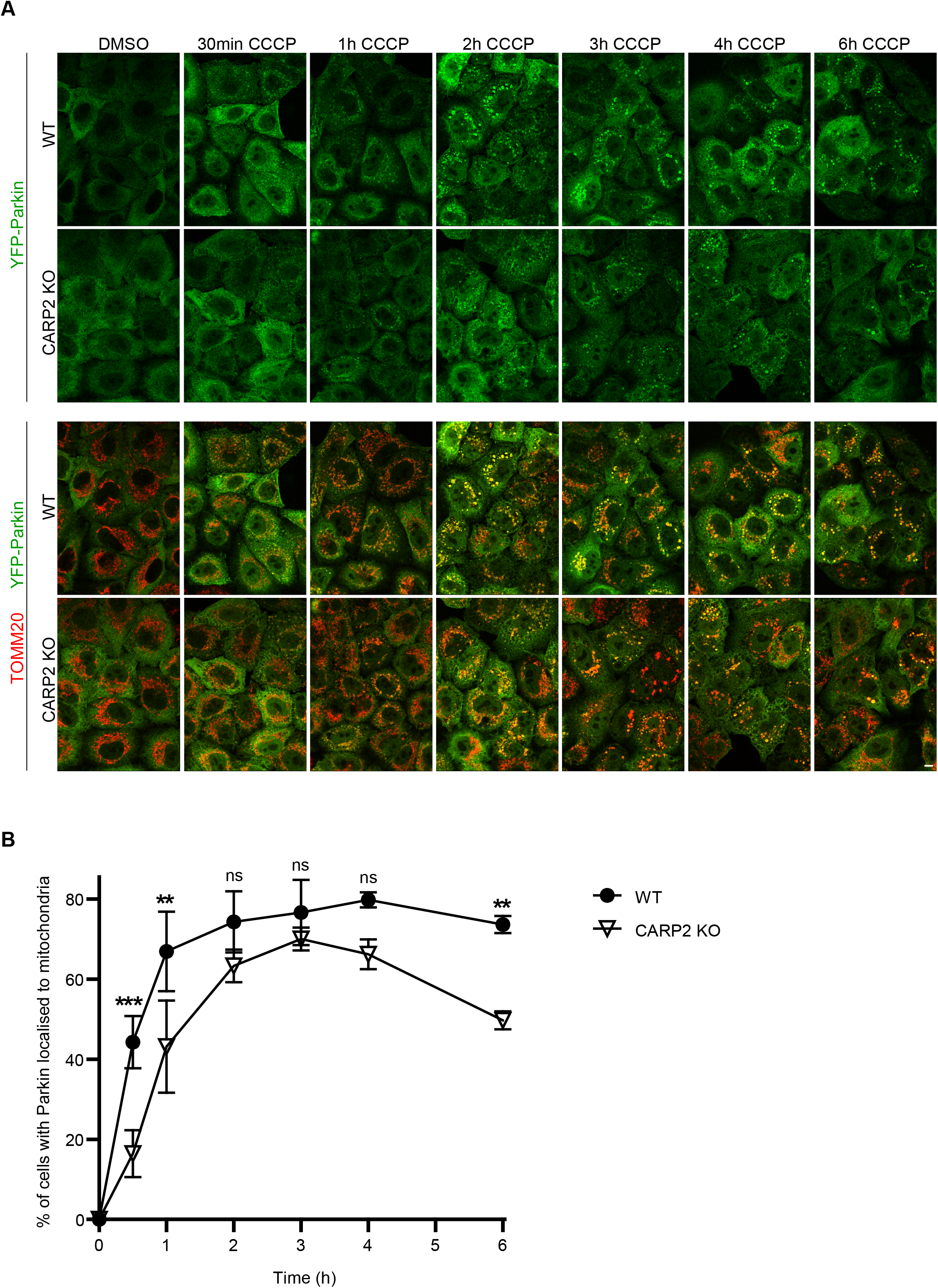
CARP2 is required for Parkin recruitment and stable association with damaged mitochondria. (**A**) A549 Wild-type or CARP2 KO cells stably expressing YFP-Parkin were treated with 10μM CCCP for 30min to 6h. Images of cells immunostained with anti-GFP and anti-TOMM20 are shown. Scale 5μm. (**B**) The graph represents the percentage of cells showing association of mitochondria with YFP-Parkin (cells having at least fifteen YFP-Parkin puncta localized to MitoTracker were considered for counting) at indicated time points. Data is from three independent experiments and a minimum of 50 cells were counted per experiment. Error bars represent mean ± SEM from three independent experiments. Statistical significance was calculated using ordinary two-way ANOVA followed by Sidak’s multiple comparisons test. P-values for C2 KO vs WT in 30min is 0.0002, 1h is 0.0039, 2h is 0.3587, 3h is 0.9417, 4h is 0.1617, and for 6h is 0.0023.

**Supplemental figure S4.**
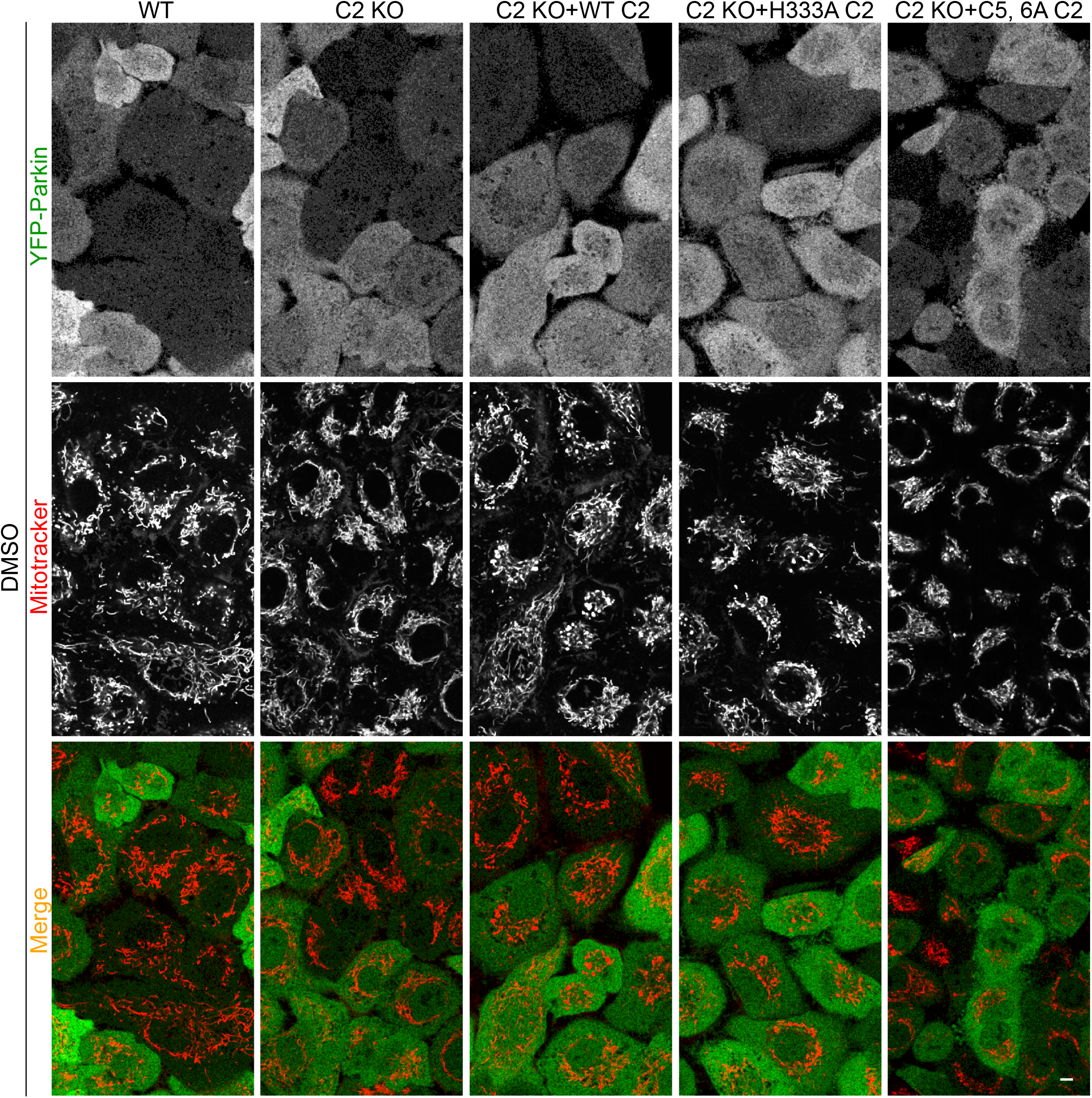
Localization of YFP-Parkin in DMSO treated A549 cells. A549 or CARP2 KO or CARP2 KO cells reconstituted with untagged CARP2 WT or H333A or C5, 6A were pre-stained with MitoTracker Red CMXRos for 30min and washed with DMEM. Subsequently, cells were treated with DMSO. Representative images of live cells are shown as a control for Figure 4C and D. Scale 5μm.

**Supplemental figure S5.**
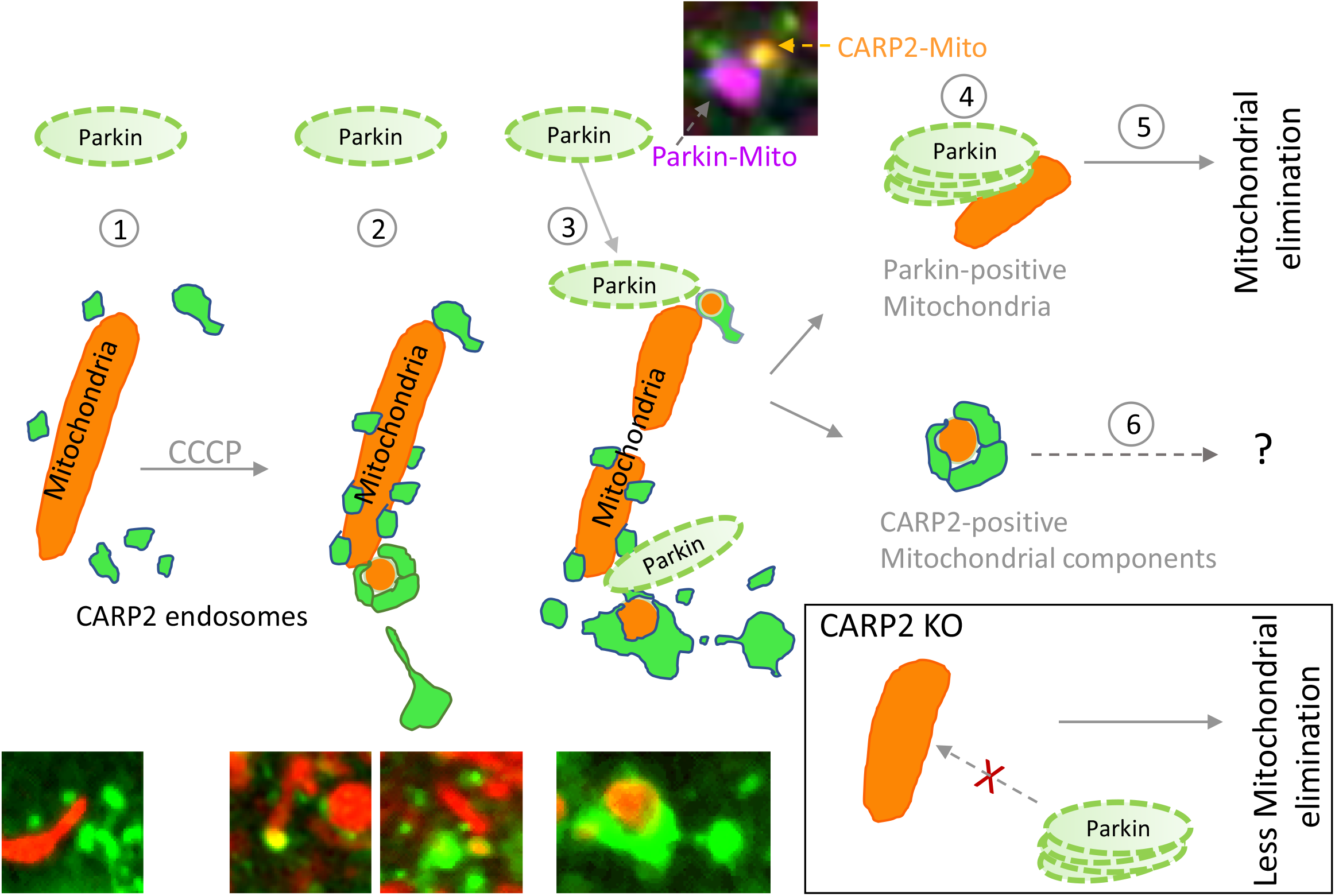
Schematic diagram outlining a working model of CARP2 in Parkin-dependent mitochondrial clearance. (1) CARP2-positive endosomal vesicles transiently associate with mitochondria. (2) Mitochondrial content during the earlier stage of damage is surrounded by CARP2 vesicles. (3) Parkin localizes with the fragmented mitochondria in close proximity with CARP2-positive mitochondria. (4) Parkin stably associates with damaged mitochondria. (5) Parkin associated mitochondria undergo mitophagy. (6) The mechanism that cells employ to eliminate CARP2-mitochondrial content preferentially either through autophagic machinery or other pathways yet to be elucidated. Confocal images of different stages of association between CARP2- and Parkin-positive mitochondria are shown. Schematics of proposed events in cells without CARP2 (CARP2 KO) are also included where Parkin stable association to damaged mitochondria are impaired, resulting in less mitochondrial elimination.

